# G4 Eurasian avian-like H1N1 swine influenza viruses exhibit enhanced pathogenicity potential in mice and pigs

**DOI:** 10.64898/2026.05.12.724537

**Authors:** Jun Jiao, Jin Ding, Ziyan Sun, Chenglin Chi, Sen Jiang, Nanhua Chen, Wanglong Zheng, Chaoyang Chen, Weiwei Su, Xiaoyan Ding, Jianzhong Zhu

## Abstract

Currently circulating swine influenza viruses (SIVs) mainly include H1N1, H1N2, and H3N2 subtypes. In this study, two G4 genotype Eurasian avian-like (EA) H1N1 SIVs were isolated from 556 samples collected between 2023 and 2026. A systematic analysis was conducted on the two EA H1N1 isolates (FYD30 and YZF69) to assess their pandemic potential. The hemagglutinin (HA) proteins of both H1N1 viruses possessed residues 225E and 228S, indicating enhanced affinity for human-like α-2,6-linked sialic acid receptors, which was confirmed by receptor-binding assays. Polymerase activity tests demonstrated that the two SIVs exhibited significantly higher activity in mammalian cells, relative to avian cells, which is consistent with the efficient replication in mammalian cells. Challenge experiments revealed that both H1N1 caused significant pathogenicity in mice and pigs, with YZF69 exhibited higher virulence than FYD30. The higher virulence of YZF69 may be attributed to its molecular features, including the NP Q357K mutation, and an additional glycosylation site in HA. In conclusion, currently circulating EA H1N1 SIVs have acquired key molecular signatures of mammalian adaptation, exhibit enhanced virulence in mammals, and continue to undergo extensive reassortment driven by international swine trade. These findings highlight the potential pandemic risk of SIVs and underscore the urgent need for strengthened surveillance.

## Introduction

Influenza A viruses pose a persistent and significant threat to global public health due to their capacity for cross-species transmission and rapid evolution. Historically, major pandemics such as the 1918 “Spanish flu”, the 1957 “Asian flu”, the 1968 “Hong Kong flu” and the 2009 H1N1 pandemic were all caused by reassortant viruses, underscoring the recurring risk of novel influenza strains emerging in human populations(1). Pigs play a particularly critical role in this dynamic: their respiratory epithelial cells express both α-2,3-linked (avian-preferred) and α-2,6-linked (human-preferred) sialic acid receptors, enabling simultaneous infections by avian and human influenza viruses(2). As a result, swine are often regarded as “mixing vessels” capable of fostering the emergence of pandemic reassortant strains(2, 3).

The 2009 H1N1 influenza pandemic (pdm/09 H1N1) served as a stark reminder of this risk. Following its emergence, the pdm/09 virus was repeatedly introduced into swine populations worldwide, where it underwent extensive reassortment with endemic strains such as classical swine H1N1 and H3N2 viruses. This process gave rise to a wide array of novel reassortants carrying pdm/09-derived genes(4, 5), many of which exhibit enhanced adaptability to humans and present serious public health concerns. Moreover, human infections caused by triple-reassortant viruses carrying pdm/09 genes are frequently reported(6, 7).

China harbors a particularly complex ecosystem of swine influenza viruses (SIVs), being the only known region where North American-lineage triple-reassortant viruses co-circulate with Eurasian avian-like (EA) H1N1 viruses(8, 9). This unique viral milieu provides an ideal environment for further reassortment. Since 2016, a specific reassortant-termed the G4 genotype has become the predominant Eurasian avian-like H1N1 strains in Chinese swine populations(10). Reports confirm that G4 EA H1N1 viruses continued to dominate in Chinese pig herds between 2019 and 2023(11). These viruses retain the HA and NA genes from Eurasian avian-like lineages, while their viral ribonucleoprotein complex and matrix protein genes are derived from the pdm/09 virus. It possesses all the hallmark features of a virus highly adapted to infect humans(10).

Against this backdrop, we conducted whole-genome sequencing and phylogenetic analysis on two isolated H1N1 swine influenza virus (SIV) strains. To further assess viral replication capacity and pathogenicity, we carried out both *in vitro* experiments and *in vivo* infections using mice and pigs. The findings reveal that internal genes originating from the pdm/09 lineage have undergone sustained and rapid evolution in swine, which significantly enhances the pathogenicity of G4 Eurasian H1N1 SIVs in mammalian hosts. This study provides crucial scientific insights into the evolutionary mechanisms and interspecies transmission potential of G4 H1N1 influenza A viruses, offering important implications for pandemic preparedness and the formulation of effective prevention and control strategies.

## Results

### Survey of clinical samples and isolation of H1N1 viruses

During the period from September 2023 to January 2026, a total of 556 samples were collected in this study. Ten of these samples tested positive and were subsequently used for virus isolation by inoculation into both MDCK cells and specific pathogen-free (SPF) chicken embryos (Table 1). Clear hemagglutination (HA) activity was detected in both the cell culture supernatants harvested 72 h post-inoculation and the allantoic fluids collected from chicken embryos 96 h after inoculation. Full-genome sequencing was performed using the HA and NA primers(12). Phylogenetic analysis, performed by comparing the HA and NA gene sequences with representative strains obtained from the NCBI database, led to the identification of two distinct H1N1 swine influenza viruses (SIVs) (Table 1). These two strains have been designated as A/Swine/Anhui/FYD30/2025 and A/Swine/Jiangsu/YZF69/2024.

**Table 1.**
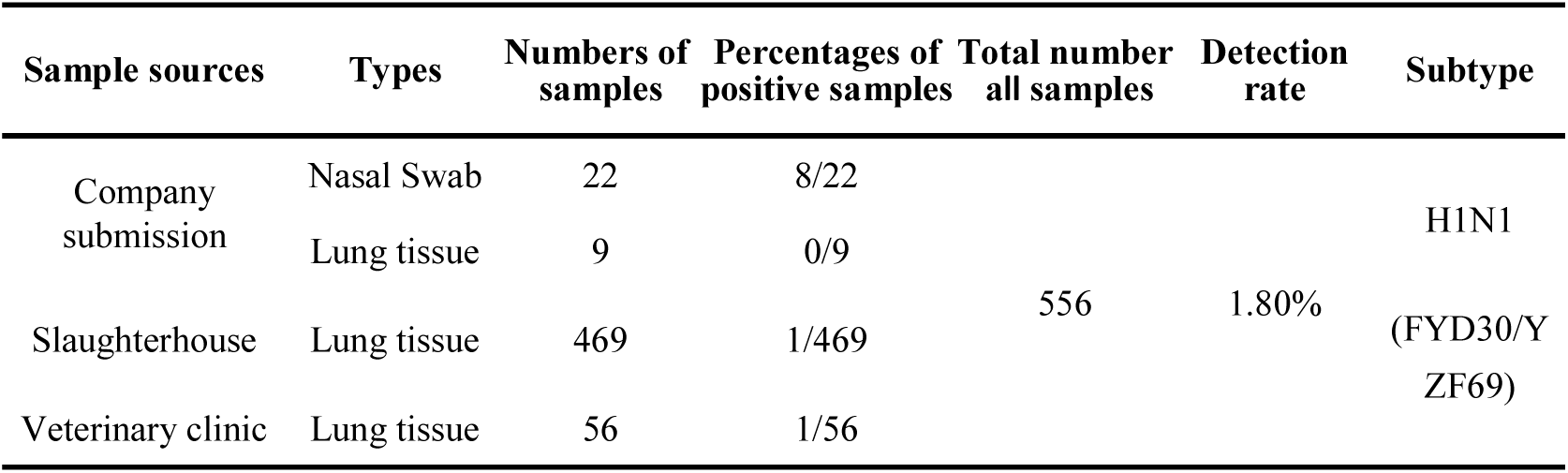
Results of swine influenza virus (SIV) detection.

### Successful rescue of two H1N1 SIVs by reverse genetics

Gene-specific primers incorporating *Bsm*BI restriction sites were used to amplify all eight gene segments of the FYD30 and YZF69 SIVs. These segments were subsequently ligated into the *Bsm*BI-linearized pHW2000 vector using seamless cloning technology. Following sequence confirmation, the plasmids were mixed in equimolar ratios and co-transfected into 293T cells. Supernatants (passage P0 stocks) were harvested 48-72 h post-transfection and inoculated into fresh MDCK cells for amplification, followed by three rounds of blind passaging. The resulting P1 viral supernatants exhibited clear hemagglutination activity, confirming the successful rescue of recombinant rFYD30 and rYZF69 viruses bearing functional hemagglutinin.

### Genetic evolution and homology analysis of two H1N1 SIVs

Phylogenetic trees for all eight gene segments were reconstructed by aligning representative sequences of the H1N1 lineage, obtained from the GISAID and NCBI databases, with the isolated strains FYD30 and YZF69 (Fig 1). Phylogenetic reconstruction revealed that the HA and NA genes of the H1N1 isolates clustered within the Eurasian avian-like lineage (Fig 1A-B). The NS gene was classified into the triple-reassortant lineage (Fig 1C), while the remaining internal genes (M, NP, PB1, PB2, PA) all grouped within the pandemic 2009 lineage (Fig 1D-H). This genetic configuration corresponds to the G4 genotype H1N1 SIVs, which is currently the predominant genotype in circulating SIVs.

**Fig 1.**
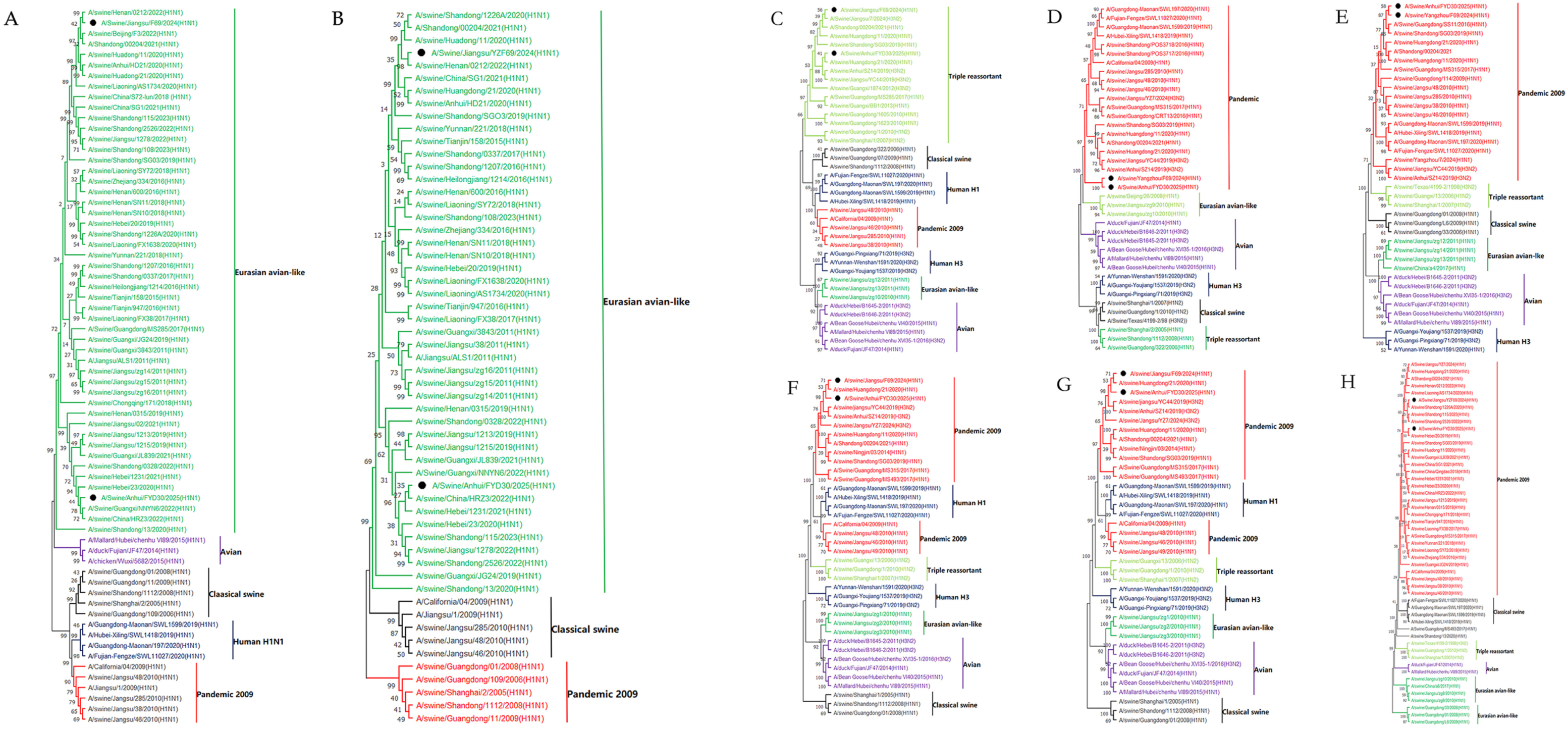
Phylogenetic trees of two isolated H1N1 swine influenza viruses. Phylogenetic analysis of the eight genes of the isolated viruses: (**A**) HA gene, (**B**) NA gene, (**C**) NS gene, (**D**) M gene, (**E**) NP gene, (**F**) PB1 gene, (**G**) PB2 gene, and (**H**) PA gene, was conducted using the neighbor-joining method in MEGA 11. The reliability of each tree node was assessed through bootstrap analysis with 1,000 replicates. The viruses isolated in this study are indicated by solid black circles.

BLAST analysis against the NCBI database was conducted to identify reference sequences with the highest homology to each gene segment of the two H1N1 isolates (Table 2). For strain FYD30, its HA and NA genes shared the greatest sequence identity with A/swine/China/SD6591/2019 (H1N1), a Shandong isolate obtained in 2019. For strain YZF69, the HA, NA, and NS genes exhibited the highest homology with A/swine/Jiangsu/HD11/2020 (H1N1), a Jiangsu isolate from 2020. In contrast, its PB1, PB2, and PA genes showed the closest match to strains circulating in Liaoning, China. (Table 2).

**Table 2.** The reference virus strains with the highest homology for each gene segments of the two H1N1 isolates.

| Strain Name | Gene | Closest Virus | Homology (%) |
| --- | --- | --- | --- |
| H1N1 (YZF69) | HA | A/swine/Jiangsu/HD11/2020(H1N1) | 97.12% |
|  | NA | A/swine/Jiangsu/HD11/2020(H1N1) | 98.02% |
|  | M | A/swine/China/SD6591/2019(H1N1) | 98.68% |
|  | NS | A/swine/Jiangsu/HD11/2020(H1N1) | 98.57% |
|  | NP | A/Singapore/276/2009(H1N1) | 96.57% |
|  | PB1 | A/swine/Liaoning/TL5404/2020(H1N1) | 96.70% |
|  | PB2 | A/swine/Liaoning/JZ164/2020(H1N1) | 96.19% |
|  | PA | A/swine/Liaoning/AS1732/2020(H1N1) | 97.46% |
| H1N1 (FYD30) | HA | A/swine/China/SD6591/2019(H1N1) | 95.56% |
|  | NA | A/swine/China/SD6591/2019(H1N1) | 97.67% |
|  | M | A/Singapore/471/2009(H1N1) | 97.10% |
|  | NS | A/swine/China/2328/2020(H1N2) | 96.30% |
|  | NP | A/swine/Guangxi/BB1/2013(H1N1) | 97.09% |
|  | PB1 | A/swine/China/SD6591/2019(H1N1) | 97.58% |
|  | PB2 | A/Hamburg/NY1580/2009 (H1N1) | 96.04% |
|  | PA | A/Sus scrofa/Jiangsu/SIV-PA/2022 (H1N1) | 95.66% |

Note. The reference isolates of HA and NA genes are blue marked, and the foreign reference isolates are green marked.

Notably, the reference SIV strains most closely related to our isolates exhibited very diverse origins of geographically regions, including different countries (Table 2). Specifically, the references SIVs with the highest M and PB2 homology to FYD30 were come from Singapore and USA, respectively, whereas the reference SIV with the highest NP homology to YZF69 was isolated from Singapore. It suggested that viral reassortment may be associated with swine trades or human activities.

### H1N1 genome molecular features analysis

Genomic analysis of two H1N1 SIV strains revealed that the HA protein of FYD30 acquired the 190D mutation, according to H3 numbering. Additionally, the presence of 225E and 228S indicated that two H1N1 have evolved a preferential binding affinity for human-like α2-6-linked sialic acid receptors (Table 3). Concurrently, multiple molecular markers associated with enhanced mammalian adaptation and virulence were also identified(13, 14). Specifically, the FYD30 and YZF69 isolates carry mutations in the PB2 protein, specifically 271A, 590S and 591R (15–17), yet lack common mammalian adaptation markers such as 627K and 701N (Table 3). These mutations significantly enhance viral replication efficiency in mammalian cells and swine hosts. Crucially, previous studies have demonstrated that the presence of 271A, 590S and 591R can functionally compensate for the absence of 627K and 701N (18). Several amino acid substitutions in the PA proteins of two H1N1 isolates, specifically 97I, 85I, 186S and 336M, have been associated with increased polymerase activity or enhanced virulence of influenza viruses (19–25) (Table 3). It is noteworthy that among the mammalian adaptive mutations identified in Eurasian H1N1 SIVs, the NP Q357K substitution serves as a key determinant of the virulence phenotype in mice (26). In this context, the FYD30 isolate carries the 357E mutation at the corresponding position, whereas YZF69 isolate harbors the 357K mutation (Table 3). The FYD30 and YZF69 isolates both carry the 215A and 30S mutations in the M1 protein (Table 3), suggesting that these viruses may exhibit enhanced virulence in a mouse model(27). Meanwhile, the stable presence of the 42S and 92D mutations in the NS1 protein confers an improved ability to evade interferon-mediated innate immune responses (28) (Table 3). All these results indicated that the SIVs not only exhibit enhanced pathogenicity in mammals and efficient transmission among swine populations, but also possess the molecular basis for infecting humans.

**Table 3.** Molecular characteristics of two isolated H1N1 SIVs.

| Genes | Functions of the residues | Residues in references | Residues in this study |
| --- | --- | --- | --- |
| **HA** | Effective binding to α2-6-linked sialic acid receptor | 190D, 225D/E, 228S(13, 14) | 190D (FYD30), 190N (YZF69), 225E (2), 228S (2) |
|  | Antigenicity affecting Eurasian avian-like H1N1 swine influenza virus | E158G(36) | 158G (2) |
| **NA** | Neuraminidase inhibitor resistance | H275Y, I223R(33-35)<br>E119V, R152K,<br>R292K(31, 32) | 275H (2), 223I (2), E119 (2), R152 (2), R292 (2) |
| **PB2** | Critical for efficient replication and adaptation of swine influenza virus in mammals | T271A, A590S, A591R(18) | 271A (FYD30) / 271S (YZF69), 590S (2), 591R (2) |
|  | Effects on the host range and virulence of influenza viruses | 627K(37), 701N(38) | 627E (2), 701D (2) |
| **PA** | Enhancement of virulence and polymerase activity in Eurasian avian-like H1N1 swine influenza virus | 100I, 321K, 330V, T97I, 85I, 186S, L336M(19-25) | 100I (2), 321K (2), 330V (2), T97I (2), 85I (2), 186S (2), L336M (2), |
| **NP** | Determination of the virulence phenotype in mice | Q357K (26) | 357E (FYD30), 357K (YZF69) |
| **NS1** | Escape from interferon-mediated innate immune responses | 42S, 92D(28) | 42S (2), 92D (2) |
| **M1** | Enhanced pathogenicity in mice | 215A, 30S(27) | 215A (2), 30S (2) |
| **M2** | Resistance to amantadine-based antivirals | L26F, V27A, A30T/V, S31N(29, 30) | 26L (2), 27V (2), 30A (2), 31N (2), 34G (2) |

Within the M2 protein, the amino acid substitutions primarily associated with amantadine resistance are L26F, V27A, A30T, and S31N. Among these, the S31N mutation plays a pivotal role in conferring drug resistance (29, 30). Notably, both isolates harbor the S31N mutation in their M2 proteins (Table 3), indicating that they have acquired resistance to amantadine. Analysis of the NA protein, however, revealed no mutations at key resistance sites, including H275Y, I223R, E119, R152, and R292 (31–35) (Table 3), indicating that the viruses remain susceptible to neuraminidase inhibitors. These findings offer valuable reference points for the clinical management of human influenza virus infections.

### Analysis of HA protein glycosylation sites and prediction of B-cell epitopes

Glycosylation sites play a pivotal role in modulating the antigenicity and receptor-binding properties of the HA protein (39). They are also crucial for viral antigenic drift and have significant implications for vaccine design (40, 41). In this study, the two isolates exhibited distinct glycosylation profiles. Specifically, strain FYD30 contains four such sites, positions 28 (NSTD), 40 (NVTV), 498 (NGTY), and 557 (NGSL), while lacking the site at position 212 (NHTY) (Table 4). In contrast, strain YZF69 harbors five potential glycosylation sites at positions 28 (NSTD), 40 (NVTV), 212 (NHTY), 498 (NGTY), and 557 (NGSL) (Table 4). To assess the antigenic differences between the two H1N1 viruses, we employed BepiPred-3.0 to predict B-cell epitopes on their HA proteins. The analysis revealed that strain FYD30 exhibits a strong B-cell epitope propensity (with scores exceeding the threshold) at both the Ca1 antigenic site (residues 170–171) and the Ca2 antigenic site (residues 200–206) (Fig 2A). In contrast, strain YZF69 showed epitope prediction only at the Ca2 site (residues 200–204), while no significant epitope signal was detected at the Ca1 site (residues 170–171) (Fig 2B).

**Fig 2.**
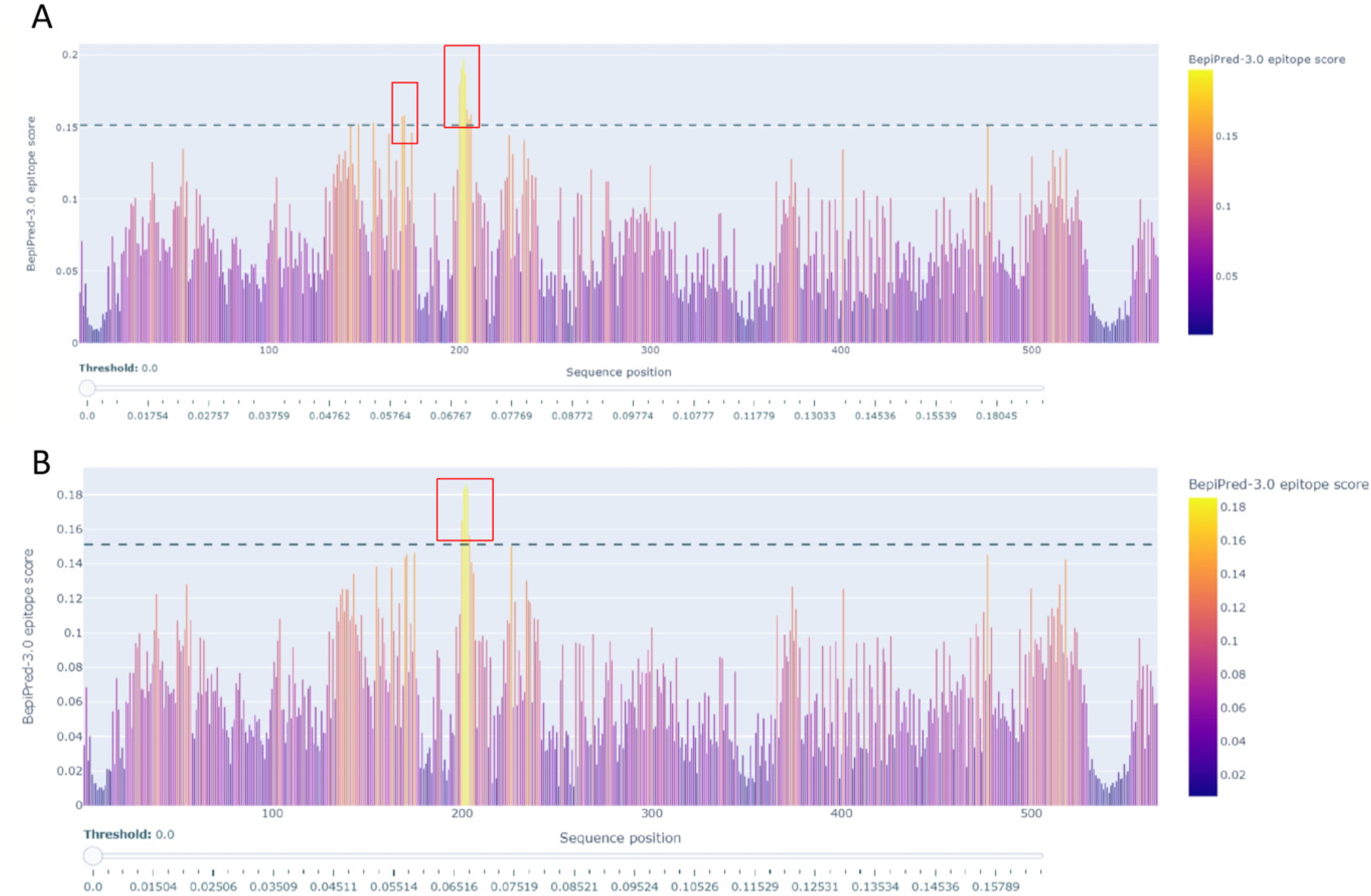
Prediction of linear B-cell epitopes in the HA proteins of FYD30 and YZF69 H1N1 swine influenza viruses. Linear B-cell epitopes in the HA protein sequences of the FYD30 **(A)** and YZF69 **(B)** strains were predicted using the BepiPred-3.0 tool (threshold = 0.5). Predicted epitope regions are indicated by red boxes. Key antigenic sites (Ca1, residues 169–173; Ca2, residues 203–219) are shown.

**Table 4.** Glycosylation sites of HA proteins of two H1N1.

| Positions | Sequence Motifs | Potential | YZF69-N-Glyc Result | FYD30-N-Glyc Result |
| --- | --- | --- | --- | --- |
| 27 | NNST | 0.4144 | – | – |
| 28 | NSTD | 0.7960 | + + + | + + + |
| 40 | NVTV | 0.7511 | + + + | + + + |
| 212 | NHTY | 0.5183 | + | - |
| 291 | NCTT | 0.4624 | - | - |
| 498 | NGTY | 0.5153 | + | + |
| 557 | NGSL | 0.6737 | + + | + + |

Note: The threshold for NetNGlyc 1.0 prediction was set at 0.5. Sites with a potential score $\geq 0.5$ and Jury agreement reaching (+) or (++) were considered potential N-glycosylation sites.

What makes this finding particularly intriguing is that the core region of the Ca1 antigenic site (residues 170-171) shares an identical amino acid sequence between the two strains, yet their epitope prediction scores differ markedly. This discrepancy suggests that local conformational changes or mutations in neighboring residues of the YZF69 HA protein may have altered the antigenic exposure of this region, potentially leading to the “masking” of epitope or a reduction in its immunogenicity.

### H1N1 replications in different cell types and polymerase activities

To evaluate the *in vitro* replication capacity of H1N1 SIVs, we measured the replication kinetics of the rFYD30 and rYZF69 strains in multiple cell lines, including MDCK cells, porcine alveolar macrophages (3D4/21), human lung carcinoma cells (A549), and chicken macrophages (HD11). The results showed that at a multiplicity of infection (MOI) of 0.001, the FYD30 strain exhibited efficient replication in MDCK cells, with a peak titer reaching 5.667-6.333 log_10_ TCID_50_/mL. Its replication in 3D4/21 cells was slightly lower, at 5.000-5.167 log_10_TCID_50_/mL, while in human A549 cells, the titer ranged from 4.000 to 4.333 log_10_ TCID_50_/mL (Fig 3A). In contrast, replication was limited in avian-origin HD11 cells, with titers only reaching 2.500-3.000 log_10_ TCID_50_/mL (Fig 3A). The replication ability of the YZF69 strain was consistently lower than that of FYD30 across all cell types tested. Peak titers for YZF69 were 5.500-6.167 log_10_ TCID_50_/mL in MDCK cells, 4.333-4.667 log_10_ TCID_50_/mL in 3D4/21 cells, 3.500-3.667 log_10_ TCID_50_/mL in A549 cells, and 3.000-3.333 log_10_ TCID_50_/mL in HD11 cells, respectively (Fig 3B).

**Fig 3.**
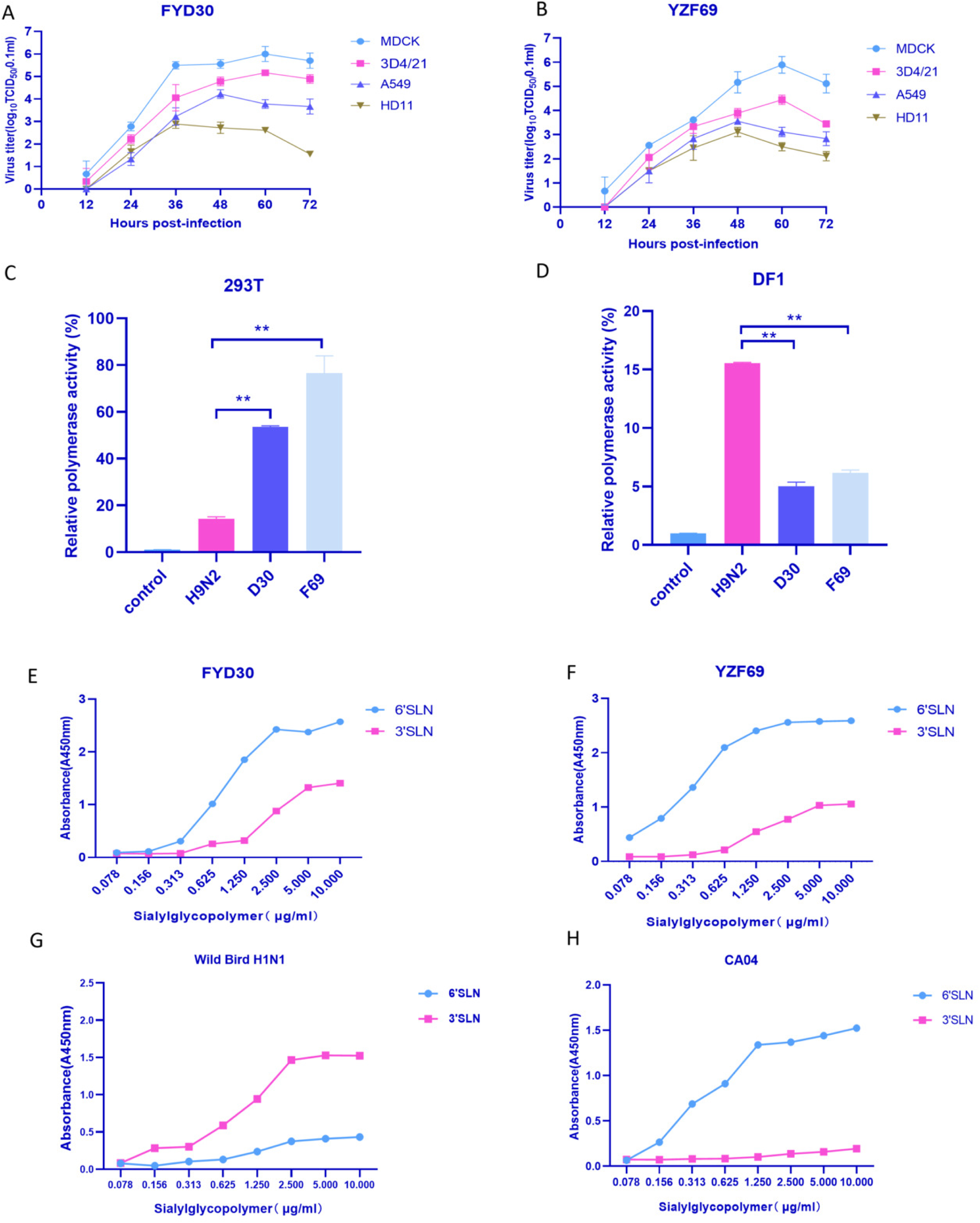
Growth kinetics, polymerase activity assay and receptor binding of two H1N1 SIVs. **(A** and **B)** MDCK, 3D4/21, A549 and HD11 cells were infected with equivalent amounts of FYD30 (A) and YZF69 (B) viruses. Cell supernatants were collected at the indicated time points post-infection and titrated for TCID_50_ in MDCK cells. **(C** and **D)** Comparison of polymerase activity of FYD30, YZF69, and avian H9N2 viruses in HEK293T cells (C) and DF-1 cells (D). Numbers represent fold change in polymerase activity compared to the control groups. (**E** and **F**) The receptor-binding properties of FYD30 (E) and YZF69 (F) were assessed using a glycopolymer-based ELISA. The avian-origin H1N1 virus **(G)** binds exclusively to 3’SLN, while the human seasonal H1N1 virus **(H)** binds exclusively to 6’SLN. All data are expressed as mean ± SD of three replicates. **p < 0.01.

Both H1N1 SIVs were thus capable of replicating in avian and mammalian cells, but viral titers in HD11 cells were markedly lower at all time points compared with those in mammalian cells. These findings indicated that FYD30 and YZF69 possess the molecular foundation for efficient replication in mammalian hosts, while exhibiting reduced adaptability to avian-derived cells.

To further investigate the underlying reason, we assessed two H1N1 polymerase activities in both avian and mammalian cells. The results showed that in mammalian 293T cells, the polymerase activities of FYD30 and YZF69 were significantly higher than that of an avian H9N2 influenza virus (Fig 3C). In contrast, in DF-1 chicken embryo fibroblast cells, FYD30 and YZF69 displayed weak polymerase activity while H9N2 exhibited higher polymerase activity in DF-1 cells (Fig 3D).

Our phylogenetic analysis showed that the PB2, PB1, PA, and NP genes (which constitute the vRNP complex) of both H1N1 SIV isolates originated from the 2009 pandemic (pdm/09) genetic lineage. Previous studies have suggested that the accumulation of multiple amino acid substitutions in pdm/09-derived vRNP genes can alter viral polymerase activity(11). The enhanced polymerase activities of two H1N1 SIVs may be attributed to cumulative mutations in their pdm/09-derived vRNP genes, which promote more efficient polymerase function in mammalian cells.

### Receptor binding specificity of the two H1N1 viruses

To evaluate the receptor-binding profiles of two H1N1, we assessed rFYD30 and rYZF69 using a glycopolymer-based ELISA. The results demonstrated that both FYD30 and YZF69 exhibited typical dual-receptor binding characteristics, capable of binding to both α-2,3-linked sialic acid receptors (3’SLN) and α-2,6-linked sialic acid receptors (6’SLN). However, the binding affinity for 6’SLN was significantly higher than that for 3’SLN in both H1N1 stains. In the glycopolymer concentration-absorbance curves, the curves corresponding to 6’SLN displayed steeper slopes and higher plateau absorbance values, directly reflecting their stronger binding capability (Fig 3E and F). These findings suggest that although FYD30 and YZF69 retain the potential to bind avian-like receptors, they have adapted more efficiently to bind human-like receptors (6’SLN). The avian-origin H1N1 virus bound exclusively to α-2,3-linked sialic acid receptors (3’SLN) (Fig 3G), showing no significant binding to α-2,6-linked sialic acid receptors (6’SLN). In contrast, the human seasonal H1N1 virus bound exclusively to 6’SLN, with no significant binding to 3’SLN (Fig 3H).

### Pathogenicity assessment of two H1N1 in the mouse model

Mice serve as an important animal model for assessing the potential human pathogenicity of influenza viruses. This study evaluated the replication kinetics and pathogenicity of two H1N1 virus strains (rFYD30 and rYZF69) in mice. Through continuous monitoring over a 14-day period, we observed that, compared to the control group, mice infected with either rFYD30 or rYZF69 began to lose weight starting from day 2 post-infection (Fig 4A). They also exhibited clinical signs such as lethargy, huddling, reduced activity, and ruffled fur. Regarding lethality, in the rFYD30-infected group, one death occurred on day 5 post-infection, and another mouse was euthanized due to weight loss exceeding 25%. The remaining three mice returned to their initial body weight by day 11 post-infection. The rYZF69-infected group showed more severe outcomes: three deaths occurred on day 5, and by day 6, the remaining two mice were euthanized as their weight loss also surpassed 25% (Fig 4B).

**Fig 4.**
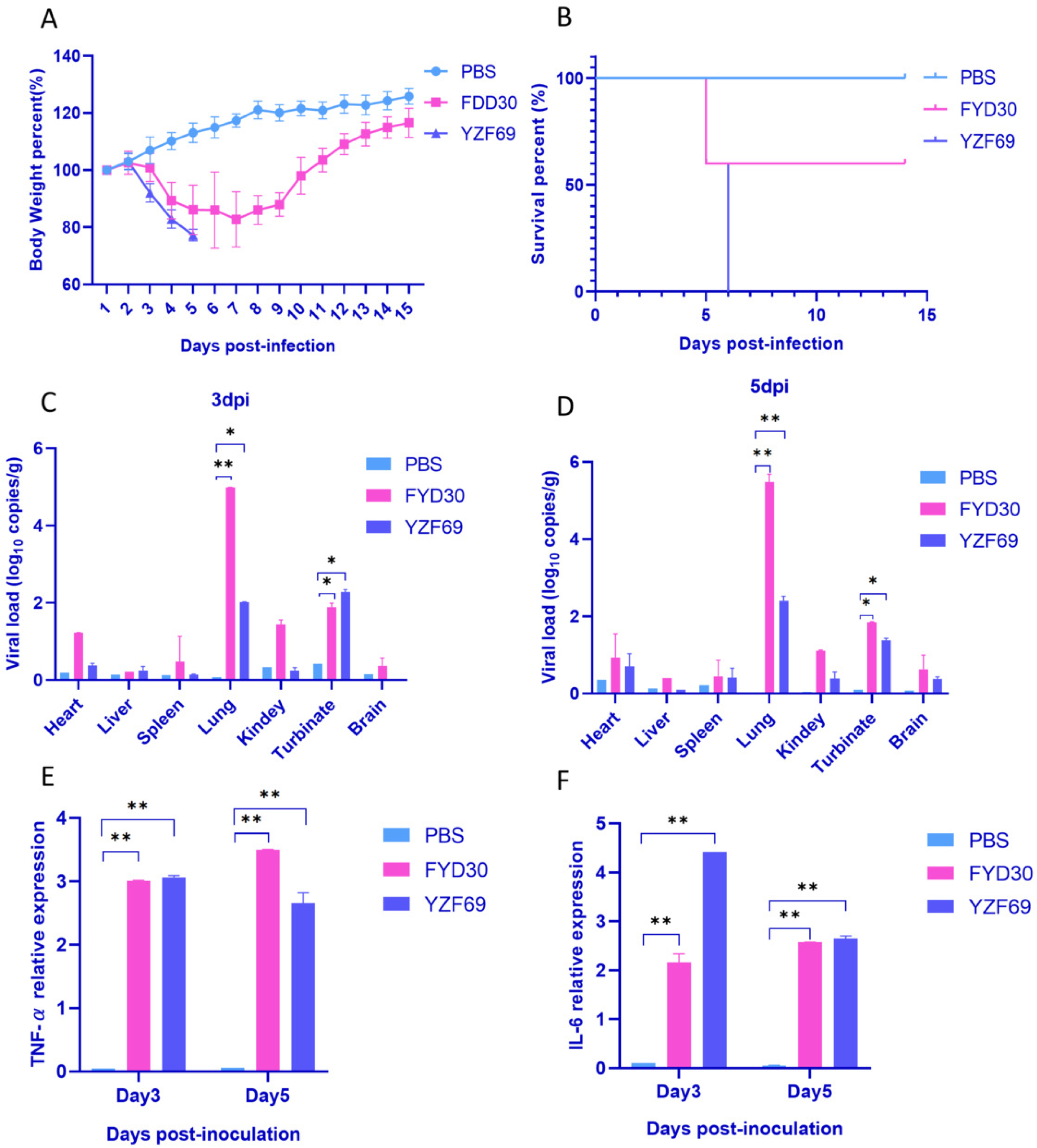
Pathogenicity of the isolated H1N1 SIVs in mice. Mice of both monitoring group and dissection group were infected with were intranasally inoculated with 10⁶ EID₅₀ of FYD30 and YZF69. Mice receiving an equal volume of PBS served as negative controls. Body weight changes **(A)** and survival rates **(B)** were monitored daily for 14 consecutive days. Viral loads in various tissues were determined by quantitative RT-PCR at day 3 **(C)** and day 5 **(D)** post-infection. Levels of TNF-α (**E**) and IL-6 **(F)** inflammatory cytokines in lung tissues were measured at day 3 and day 5 post-infection. Data are expressed as mean ± SD of three replicates. * p < 0.05 and ** p < 0.01.

Quantitative RT-PCR analysis revealed that viral loads in lung tissue and nasal turbinates were significantly higher than in other examined organs on both day 3 and day 5 post-infection (Fig 4C and D). Concurrently, the levels of the pro-inflammatory cytokines TNF-α and IL-6 in the lung tissues of infected mice were significantly upregulated on both days 3 and 5 (Fig 4E and F).

Histopathological examination (H&E staining) of lung tissues further revealed substantial damage caused by the infection. On both days 3 and 5 post-infection, both groups exhibited typical lesions, including alveolar septal thickening, macrophage infiltration, and focal hemorrhage, consistent with the features of moderate viral pneumonia (Fig 5A-I). No pathological changes were observed in the negative control group (Fig 5A, D, G). Notably, a clear difference in the distribution of lesions was observed: pathological damage in the rFYD30-infected group showed a gradient pattern along the main bronchi, with more severe injury closer to the bronchi (Fig 5B, E, H). In contrast, the rYZF69-infected group exhibited more extensive and diffuse pathological changes (Fig 5C, F, I). This distinct pattern of lesion distribution may be a key reason for the stronger pathogenicity exhibited by the rYZF69 strain in mice.

**Fig 5.**
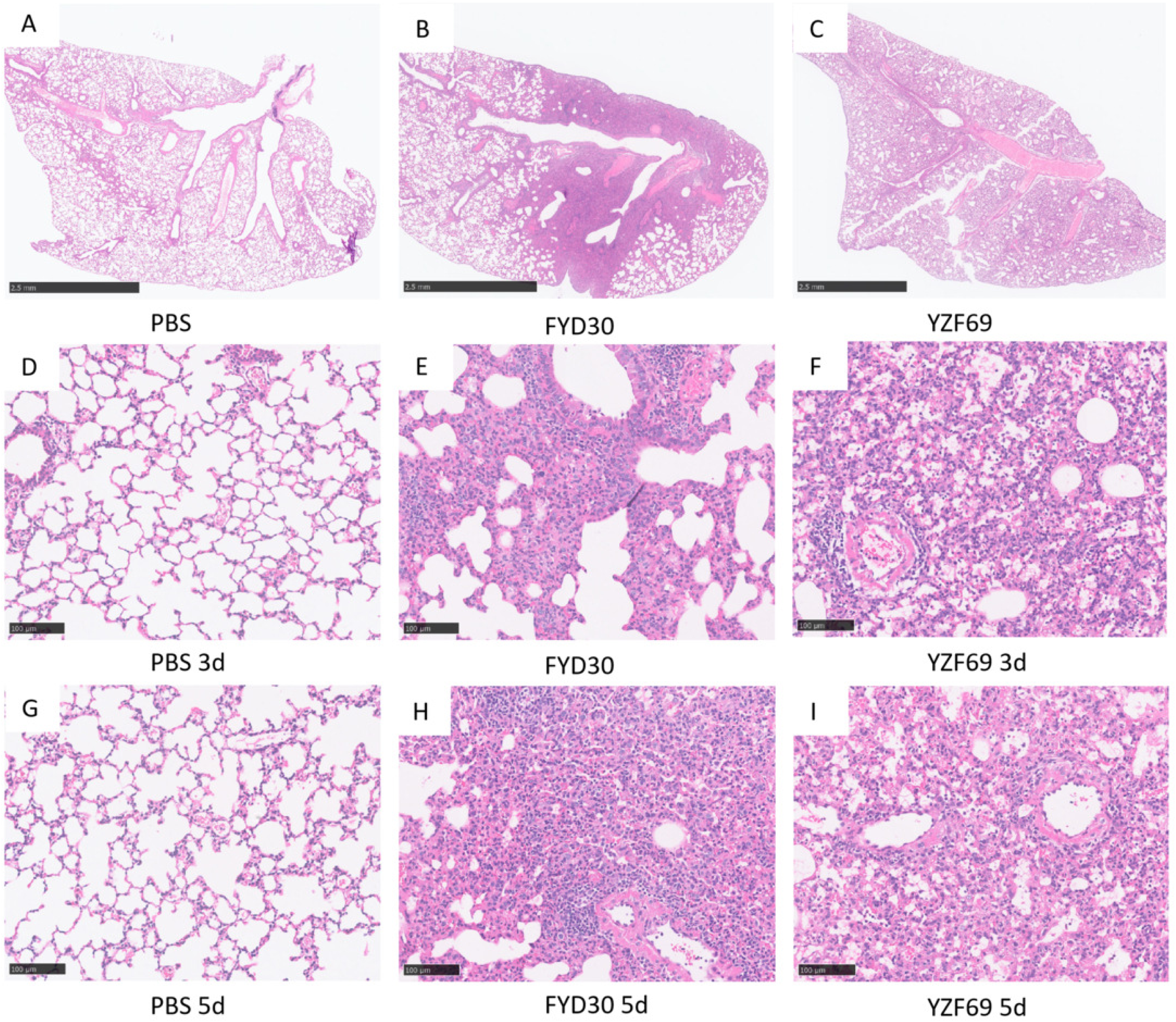
Histopathological analysis of mouse lung tissues post-infection. Mice were euthanized at 3 and 5 days post-infection, with PBS-injected mice serving as controls. Lung tissues were fixed in formalin and subjected to hematoxylin and eosin (H&E) staining for observation by microscopy. The representative sections of lung tissues from **(A)** PBS control group, **(B)** FYD30-challenged group, and **(C)** YZF69-challenged group were presented. Scale bar = 2.5 mm in panels A-C, Scale bar = 100 μm in panels D-I.

### Pathogenicity assessment of two H1N1 viruses in pigs

Following inoculation of piglets, no mortality was observed during the experiment. Compared to the PBS control group, piglets in both the rFYD30- and rYZF69-infected groups exhibited transient, mild clinical signs, primarily including sneezing, nasal discharge, and elevated body temperature. These signs were most pronounced between day 3 and day 7 post-inoculation (dpi). Temperature monitoring confirmed a febrile response in both infected groups. The peak body temperatures reached 40.5 ℃ and 40.9 ℃ for the rFYD30 and rYZF69 groups, occurring on day 5 and day 4 post-inoculation, respectively, whereas the control group maintained a normal temperature range throughout (Fig 6A). Serum antibody responses were evaluated by hemagglutination inhibition (HI) assay. Detectable HI antibodies (titer <1:16) were present in all pigs from both infected groups by 7 dpi. The antibody titers had increased significantly, reaching a maximum of 1:128 in both the rFYD30 and rYZF69 groups by 14 dpi, confirming effective infection and immune response (Fig 6B). In the meantime, serum HI antibodies remained negative in the control group throughout the study (Fig 6B).

**Fig 6.**
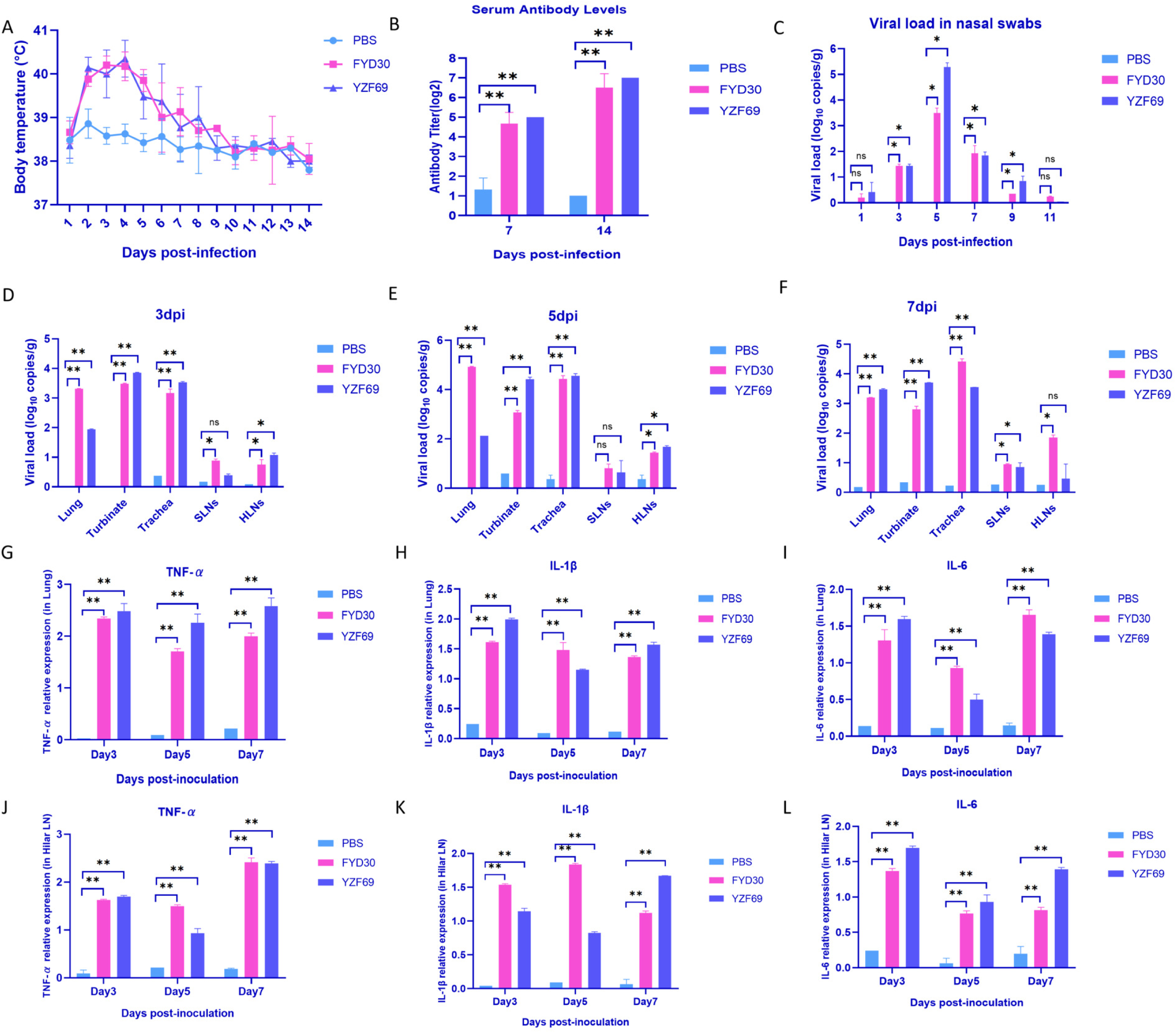
Pathogenicity of the isolated H1N1 SIVs in pigs. Pigs were intranasally inoculated with 1 mL of 10⁶ EID₅₀ of FYD30 and YZF69 viruses, with pigs receiving an equal volume of PBS as negative controls. Body weight changes were monitored till day 14 post infection **(A)**, and serum antibody levels were measured by hemagglutination inhibition (HI) assay at 7 and 14 days post-infection **(B)**. Viral loads in nasal swabs collected at 1, 3, 5, 7, 9, and 11 days post-infection were determined by quantitative RT-PCR **(C)**. Viral titers in lung tissues, nasal turbinates, trachea, submandibular lymph nodes, and hilar lymph nodes were measured at 3 days **(D)**, 5 days **(E)**, and 7 days **(F)** post-infection by quantitative RT-PCR. Levels of TNF-α, IL-1β, and IL-6 inflammatory cytokines in lung tissues **(G-I)** and hilar lymph nodes **(J-L)** were measured at 3, 5, and 7 days post-infection by quantitative RT-PCR. Data are expressed as mean ± SD of three replicates. *p < 0.05; **p < 0.01; ns, not significantly different.

Viral shedding was assessed by quantitative analysis of viral RNA in nasal swabs using RT-qPCR. Both rFYD30 and rYZF69 strains replicated efficiently, with viral shedding from the upper respiratory tract beginning at 3 days post-infection (dpi). The peak period of viral shedding for both the rFYD30 and rYZF69 groups was from 3 to 9 dpi, with peak titers of 3.77 log_10_ copies/mL for the rFYD30 group and 5.41 log_10_ copies/mL for the rYZF69 group (Fig 6C). Viral shedding in all infected animals fell below the detection limit after 11 dpi (Fig 6C). To investigate systemic viral replication, pigs were euthanized at 3, 5, and 7 dpi for viral load quantification in various tissues. The highest viral loads in both the rFYD30 and rYZF69 groups were detected in lung tissue, nasal turbinates and trachea. Notably, viral RNA was also detected in the hilar lymph nodes of both infected groups, indicating lymphotropic potential. All tissue samples from the PBS control group were negative (Fig 6D-F).

### Inflammatory response and histopathological analysis

Analysis of inflammatory cytokines (including TNF-α, IL-1β, and IL-6) in lung tissue and hilar lymph nodes revealed significant upregulation in both the FYD30 and YZF69 infected groups compared to the control group at 3, 5, and 7 days post-inoculation (dpi) (Fig 6G-L).

Regarding pathological changes, macroscopic examination of lungs from infected pigs showed noticeable alterations compared to the PBS group, including congestion, reddening, increased lung volume, and localized hemorrhagic spots (Fig 7A-I). Histopathological evaluation by H&E staining further demonstrated varying degrees of interstitial pneumonia in the lungs of both virus-infected groups. These lesions were characterized by alveolar septal thickening and inflammatory cell infiltration. The severity of pathological changes was greater in the YZF69-infected group than in the FYD30-infected group (Figs 7J-R).

**Fig 7.**
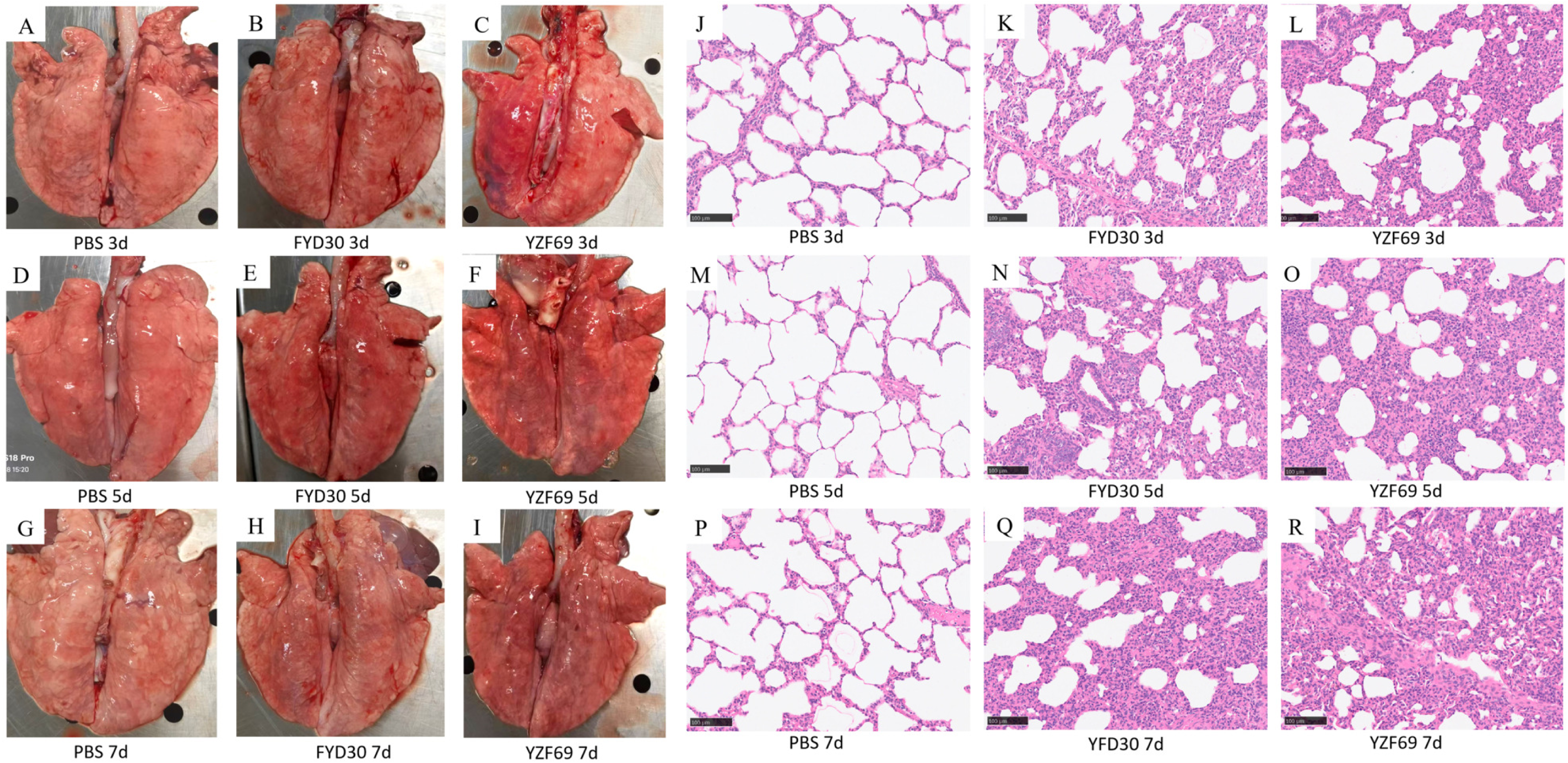
Gross lesions and histopathological changes in the lungs of infected pigs. One pig from FYD30 and YZF69 infected pigs and PBS inoculated control group was necropsied at day 3 (**A-C**), day 5 (**D-F**), and day 7 (**G-I**) post-infection, and the lung images of these pigs are shown. Lung tissues were fixed in formalin and subjected to hematoxylin and eosin (H&E) staining for observation by microscopy. The histochemical images from lung tissues of day 3 (**J-L**), day 5 (**M-O**), and day 7 (**P-R**) post-infection are presented. Scale bar = 100 μm in panels J-R.

Immunohistochemical (IHC) staining successfully detected influenza virus nucleoprotein (NP) antigen within the lung tissues of infected animals. Both the area and intensity of the NP-positive signal were greater in the YZF69 group compared to the FYD30 group (Fig 8). This co-localization of viral antigen with pathological lesions provides further evidence of active viral replication in the lung tissue and its association with tissue damage.

**Fig 8.**
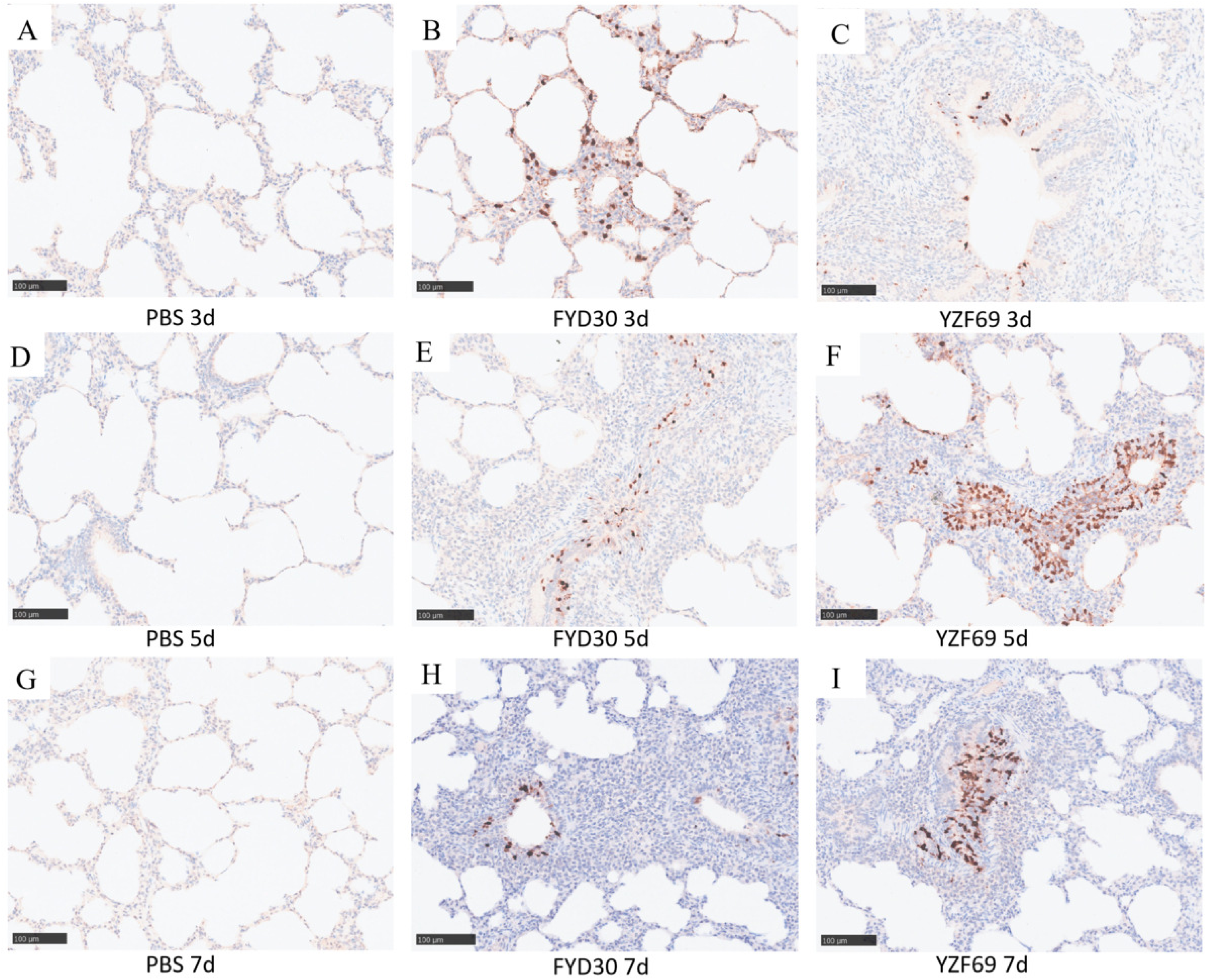
Immunohistochemical analysis of porcine lung tissues post-infection. Lung tissues were collected from each infected group and PBS control group at day 3 (**A**-**C**), day 5 (**D**-**F**), and day 7 (**G**-**I**) post-infection, fixed in formalin, and subjected to immunohistochemical staining using anti-NP antibody to observe the distribution and localization of viral antigens under microscopy. Brown staining areas indicate NP-positive signals, representing the distribution of viral antigens in lung tissues. Scale bar = 100 μm.

In summary, both the FYD30 and YZF69 strains induced a typical mild influenza infection in pigs, characterized by fever, viral shedding, specific antibody production, and pulmonary inflammation. Compared to FYD30, YZF69 demonstrated a greater capacity to induce more severe pulmonary inflammation and histopathological damage, indicating its stronger pathogenic potential pigs.

### Cross-hemagglutination inhibition assay and antigenic analysis

The cross-hemagglutination inhibition assay results showed that the antigenic correlation coefficients (R) among the three H1N1 SIVs FYD30, YZF69, and QD (isolated in 2018) were as follows: 0.86 between FYD30 and QD, 0.78 between YZF69 and QD, and 0.72 between FYD30 and YZF69 (Table 5). According to the classical criterion (*R* ≥ 0.67 indicates the same antigenic group), the three H1N1 SIV strains still fell within the same antigenic group, although the *R* value of 0.72 between FYD30 and YZF69 was close to the threshold for antigenic difference. Furthermore, the *R* values between the two H1N1 viruses and the H3N2 virus ranged from 0.47 to 0.67, with the lowest value (0.47) observed between YZF69 and H3N2, indicating a significant antigenic difference between these two strains (Table 5).

**Table 5.** Antigenic correlation coefficients (R) among different SIV strains.

| strains | R value |  |  |  |
| --- | --- | --- | --- | --- |
|  | FYD30 | YZF69 | QD | YZ07 |
|  | (2025 H1N1) | (2024 H1N1) | (2018 H1N1) | (2024 H3N2) |
| D30 | 1.00 |  |  |  |
| F69 | 0.72 | 1.00 |  |  |
| QD1 | 0.86 | 0.78 | 1.00 |  |
| H3N2 | 0.62 | 0.47 | 0.67 | 1.00 |

Note: Values represent antigenic correlation coefficients ( $R$ ). $R \geq 0.67$ indicates that the two strains belong to the same antigenic group, while $R < 0.67$ indicates a significant antigenic difference.

## Discussion

Currently, EA H1N1 SIVs have emerged in multiple countries and regions(42). Through long-term evolution and transmission, these viruses have continuously reassorted with other influenza viruses, including the 2009 pandemic H1N1 and triple-reassortant H1N2 (TR H1N2) influenza viruses, resulting in 11 genotypes to date. They have now become the predominant strains among SIVs circulating in pig populations globally(43). Studies have shown that viruses carrying the pdm/09 gene segment exhibit enhanced infectivity in humans(44). Alarmingly, seropositivity rates for antibodies against EA H1N1 have been rising in human populations, including among swine workers(45, 46). These findings constitute a clear public health alert: we must regard EA H1N1 SIV as a potential pandemic candidate and urgently strengthen surveillance and risk assessment efforts.

The two H1N1 isolates obtained in this study (FYD30 and YZF69) exhibit mammalian adaptation across multiple key dimensions. First, the presence of residues 225E and 228S in the HA protein serves as a definitive molecular signature of a receptor-binding switch from avian-type receptor preference to strong affinity for human-like α-2,6-linked sialic acid receptors abundantly distributed in the human upper respiratory tract(13, 14). Receptor-binding assays functionally validated this marked preference for human-type receptors. Consequently, both isolates have acquired the capacity to effectively attach to and invade human respiratory epithelial cells, a fundamental prerequisite for efficient human-to-human transmission. Second, polymerase activity assays revealed significantly higher replication efficiency in mammalian cells than in avian cells, corroborated by *in vitro* replication kinetics. An efficiently functioning polymerase system implies that should these viruses breach the species barrier and infect humans, they could rapidly achieve high viral loads, thereby, increasing transmission likelihood and potentially exacerbating clinical disease severity.

Phylogenetic analysis uncovered a complex genetic architecture: all five internal genes of these two isolates originate from the pdm/09 lineage, a lineage already proven capable of efficient human infection. Notably, the surface genes of strain YZF69 share high homology with A/swine/Jiangsu/HD11/2020 (H1N1), a Jiangsu isolate from 2020(47). Even more compellingly, the PA and NS genes of a previously isolated H3N2 SIV from our laboratory exhibit extremely high homology with those of the same Jiangsu isolate (48), providing direct evidence of cross-subtype reassortment among SIVs. These findings confirm that EA H1N1 viruses are not evolving in isolation; rather, they actively function as “genetic donors,” continuously exchanging gene segments with other subtypes and generating novel reassortants with complex genetic backgrounds and unpredictable biological properties. The high homology between the A/swine/Jiangsu/HD11/2020 (H1N1) isolate and human H1N1 viruses further blurs the boundary between swine and human influenza ecosystems, suggesting the possibility of bidirectional and potentially undetected gene flow between pigs and humans. Moreover, the close genetic relatedness of certain gene segments from YZF69 and FYD30 to isolates from Singapore underscores the role of international live swine trade in facilitating transboundary viral movement, serving as a stark reminder that SIVs are actively reassorting across national borders, hitchhiking with the global trade of pigs.

Concurrent animal model infection experiments demonstrated that both H1N1 SIV isolates exhibit marked pathogenicity in mice and pigs. Notably, strain YZF69 displayed significantly higher virulence in mice compared to strain FYD30, prompting an in-depth genomic sequence analysis to uncover the underlying molecular mechanisms. Sequence analysis pinpointed a key distinction: the NP Q357K substitution, established as a critical determinant of the murine virulence phenotype in Eurasian H1N1 SIVs, is present exclusively in the YZF69 isolate. Concurrently, glycosylation site analysis revealed that YZF69 possesses one additional potential N-linked glycosylation site relative to FYD30, and B-cell epitope predictions further indicated that YZF69 harbors fewer antigenic sites than FYD30. This convergence of molecular evidence strongly suggests that YZF69 is better equipped for antigenic concealment and immune evasion, mechanisms that likely synergistically contribute to its enhanced pathogenicity in mice. Although FYD30 exhibited higher viral loads in certain tissues and organs at specific time points, it induced milder pathological damage, which may be associated with differences in virus-induced apoptosis capacity, epitope masking leading to immune evasion, and other factors.

Although both FYD30 and YZF69 exhibited a preference for human-like receptors, cross-hemagglutination inhibition assays revealed that their antigenic correlation coefficient (*R*) was only 0.72, suggesting a detectable antigenic difference between the two strains. Combined with *BepiPred-3.0* epitope prediction results, we speculate that the YZF69 strain shows a loss of epitope signal at the Ca1 antigenic site (residues 170-171), possibly through conformational ‘masking’ of this epitope, thereby reducing the hemagglutination inhibition capacity of the FYD30 immune serum. Moreover, the *R* value between YZF69 and the H3N2 virus was as low as 0.47, significantly lower than that between FYD30 and H3N2 (0.67), further confirming the extent of antigenic variation in YZF69. These findings suggest that during the evolution of EA H1N1 SIVs, antigenic variation may occur prior to changes in receptor-binding properties, and certain lineages (such as YZF69) may evolve at a faster rate than others.

In summary, this study demonstrates that currently circulating EA H1N1 SIVs have acquired key mammalian adaptive markers and possess the potential to infect and transmit among mammals. Of particular concern, the continuous reassortment and antigenic variation occurring within EA H1N1 viruses, coupled with their enhanced preference for human-like receptors, further increase the risk of cross-species transmission. These findings underscore the urgency of conducting sustained, systematic surveillance and risk assessment of SIVs, especially the EA H1N1 lineage, and provide important scientific evidence for influenza pandemic preparedness and the rational selection of vaccine strains.

## Materials and methods

### Animal ethics statement

Specific-pathogen-free (SPF) BALB/c mice were purchased from Animal Facility Center, Yangzhou University, and experimental pigs and SPF chickens were purchased from Jiangsu Lihua Animal Husbandry Co., Ltd. The experiments of mice, pigs and chickens were carried out according to the laboratory animal welfare and ethical guidelines of Yangzhou University, under the certificate numbers SYXK(JS) 2022-0044, SYXK(JS) 2026-0023, SYXK(JS) 2026-0024, respectively.

### Reagents, cells and viruses

Human embryonic kidney 293T cells (HEK-293T) and Madin-Darby canine kidney (MDCK, ATCC CRL-34) cells were maintained in DMEM (Hyclone Laboratories, USA) supplemented with 10% fetal bovine serum (FBS, Vazyme Biotech, China). Porcine alveolar macrophage cell line (3D4/21) and chicken macrophage HD11 cells (BCRJ-Linhagens celulares Cat# 0099) were cultured in RPMI 1640 medium (Hyclone Laboratories) with 10% FBS. Chicken fibroblast DF-1 cells (ATCC Cat# CRL-12203) were cultured in M199-F12 (Sigma, USA) with 10% FBS. The 2×MultiF Seamless Assembly Mix was from Abclonal (Wuhan, China). *Bsm*BⅠ was purchased from NEB (Beijing, China). TPCK-trypsin was from Sigma. Two swine influenza viruses (SIVs) (FYD30 and YZF69) were isolated in this study, one SIV (QD) isolated in 2018 and an avian H9N2 influenza virus were preserved in our laboratory, and the remaining two viruses (wild bird H1N1 and human influenza virus CA04) as controls were kindly provided by Dr. Zhimin Wan of Veterinary Medicine School of Yangzhou University.

### Sample collection, detection and virus isolation

From September 2023 to January 2026, lung tissue and nasal swab samples were collected from pig farms, slaughterhouses, and veterinary clinics in northern Jiangsu Province. The samples were homogenized in DMEM basal medium supplemented with penicillin (100 U/mL) and streptomycin (100 U/mL). After centrifugation at 3000×g for 5 min, the supernatant was collected for total RNA extraction using TRIzol reagent (AG Accurate Biology, Hunan, China). The extracted RNA was reverse-transcribed into cDNA using the influenza virus-specific primer Uni12(48). Subsequently, polymerase chain reaction (PCR) was performed with conserved primers targeting the swine influenza virus (SIV) NP and M genes(48).

PCR-positive samples were sterilized by filtration through a 0.22μm membrane and inoculated into both confluent MDCK cell monolayers and 10-day-old SPF embryonated chicken eggs (Boehringer Ingelheim Weitong Biotech, Nantong, China). MDCK cells were maintained in serum-free medium containing 1 μg/mL TPCK-trypsin and cultured at 37℃ under 5% CO_2_ for 72 h. Chicken embryos were incubated at 36℃ for 96 h. After incubation, cell culture supernatants and allantoic fluids were harvested, and viral propagation was assessed by hemagglutination assay.

### Rescue of H1N1 swine influenza virus by reverse genetics

The H1N1 swine influenza virus (SIV) was rescued using a plasmid-based reverse genetics system, with primers and methodology adapted from reference(12). All eight viral gene segments were amplified by RT-PCR. The purified amplicons were individually inserted into *Bsm*BI-linearized pHW2000 vectors via a seamless cloning strategy. The ligation products were transformed into DH5α competent cells, and positive clones were selected by colony PCR followed by confirmation through DNA sequencing. An equimolar mixture of the eight verified plasmids was co-transfected into 293T cells. At 6 hours post-transfection, the medium was replaced with Opti-MEM maintenance medium containing TPCK-trypsin. Distinct cytopathic effects were typically observed within 48-72 h. The cell cultures were harvested, subjected to freeze-thaw cycles, and clarified by centrifugation to obtain the P0 virus stock. The P0 virus was subsequently amplified in MDCK cells. After 72 h incubation, the P1 virus supernatant was collected and tested by hemagglutination assay to confirm the successful rescue of infectious recombinant viruses.

### Phylogenetic analysis and viral molecular feature analysis

Viral sequencing data were assembled and curated using the SeqMan module of the Lasergene software suite (DNASTAR, USA). Reference genomic sequences were obtained from the NCBI Influenza Virus Database and the GISAID platform. Phylogenetic trees were constructed in MEGA11 using the neighbor-joining method, with the robustness of tree topology assessed by bootstrap analysis with 1,000 replicates. Based on the deduced amino acid sequences of two H1N1 SIV genomes, alignment and homology analysis were performed against various reference strains using the MegAlign module in Lasergene to characterize molecular features of viral proteins.

### Analysis of HA protein glycosylation sites

The amino acid sequences of the HA proteins were extracted from the complete viral genome sequences and submitted in FASTA format to the online server NetNGlyc 1.0 (Center for Biological Sequence Analysis, Technical University of Denmark; website: https://services.healthtech.dtu.dk/services/NetNGlyc-1.0/). This server predicts potential N-glycosylation sites based on an artificial neural network algorithm by analyzing the Asn-Xaa-Ser/Thr sequon (where Xaa is not proline) and its surrounding sequence environment. Predictions were performed using the default threshold (0.5). A site was considered a potential N-glycosylation site when the potential score exceeded 0.5 and the Jury agreement reached or exceeded (+/++). Additionally, SignalP-5.0 was used to predict signal peptides to exclude interference from glycosylation sites located in non-extracellular regions.

### Prediction of B-cell epitopes

To analyze the antigenic differences between the FYD30 and YZF69 H1N1 SIV strains, bioinformatics methods were employed to predict linear B-cell epitopes of their HA proteins. The amino acid sequences of HA proteins from both strains were submitted in FASTA format to the BepiPred-3.0 prediction tool on the IEDB online server (http://tools.iedb.org/bcell/). BepiPred-3.0 predicts linear B-cell epitopes by integrating amino acid sequence features with known epitope data. Predictions were performed using the software’s default parameters, with a threshold set at 0.5. Continuous amino acid residue regions with scores above the threshold were defined as potential linear B-cell epitopes. The prediction results were output in both graphical and tabular formats, recording the epitope propensity score for each residue.

### Receptor-binding assay

The 3 ‘SLN and 6 ’SLN sialylglycopolymer chains (Glycotech Inc. Gaithersburg, USA) were diluted to 10 μg/mL, followed by two-fold serial dilution to concentrations of 5 μg/mL, 2.5 μg/mL, 1.25 μg/mL, 0.625 μg/mL, 0.313 μg/mL, 0.156 μg/mL and 0.078 μg/mL. These diluted solutions were coated onto high-binding affinity ELISA plates (Thermo Fisher Scientific, Shanghai, China) and incubated overnight at 4 ℃, with a PBS group serving as the negative control. After washing with PBST, the plates were blocked with 5 % BSA for 8 h. Subsequently, virus samples diluted to 2^6^ hemagglutination (HA) units were added to the wells and incubated at 4 ℃ for 8 h. Following another wash with PBST, 2^8^ hemagglutination inhibition (HI) units of antiviral serum were added and incubated at 4℃ for 2 h, followed by incubation with HRP-labeled goat anti-mouse secondary antibody (4℃, 2h). Finally, color development was performed using TMB substrate for 10 min, and the reaction was terminated by adding 2 M H_2_SO_4_. The absorbance was measured at 450 nm using a microplate reader, and the receptor-binding characteristics were analyzed by constructing a glycopolymer concentration-absorbance curve.

### *In vitro* replication kinetics of H1N1 SIVs

To evaluate the replication capacity of two H1N1 SIV *in vitro*, infections were carried out in different cell types, including MDCK (Madin-Darby Canine Kidney cells), porcine 3D4/21 (porcine alveolar macrophage 3D4/21 cells), A549 (human lung carcinoma A549 cells) and HD11 (chicken macrophage cells). Cells were infected at a multiplicity of infection (MOI) of 0.001, washed three times with PBS following a 2 h adsorption period, and incubated with DMEM maintenance medium supplemented with 1 μg/mL TPCK-trypsin, at 37°C under 5% CO₂. The culture supernatants were collected at 12, 24, 48, 60 and 72 h post-infection (hpi) and stored at −80°C. Viral titers in supernatants were determined by titration using MDCK cells, and the 50% tissue culture infectious dose (TCID_50_) was calculated using the Reed–Muench method(49).

### Polymerase activity assay

Viral polymerase activity was determined using a luciferase reporter assay as previously described (11). Briefly, the PB2, PB1, NP, and PA genes from two H1N1 SIV strains were individually cloned into the pEGFP-C1, pCAGGS-2HA, pHW2000, and pCAGGS-3×Flag vectors, respectively. DF-1 cells or HEK293T cells were co-transfected with PB2, PB1, NP, and PA expression plasmids (100 ng each), together with the firefly luciferase reporter plasmid p-Luci (10 ng) and the internal control plasmid pRL-TK (2.0 ng). At 24 h post-transfection, cell lysates were prepared and luciferase activities were measured using the Dual-Luciferase Reporter Assay System (Vazyme Biotech, Nanjing, China) and a 96-well microplate luminometer.

### Pathogenicity assessment in a mouse model

Twenty-seven 4-to 6-week-old specific pathogen-free (SPF) female BALB/c mice were randomly divided into three groups: a dissection group (6 mice each for rFYD30 and rYZF69), a weight monitoring group (5 mice each for rFYD30 and rYZF69), and a negative control group (5 mice). Under anesthesia, mice in the dissection and weight monitoring groups were inoculated intranasally with 10^6^ EID_50_ of the rescued rFYD30 and rYZF69 influenza virus strains, respectively, while the negative control group received an equivalent volume of PBS.

Mice in the dissection groups were euthanized at 3 and 5 d post-infection (3 mice per time point per group), and one mouse in the control group was euthanized at each corresponding time point. Tissue samples including heart, liver, spleen, lung, kidney, brain, and nasal turbinates were collected for SYBR green quantitative RT-PCR analysis of virus loads (Table 6). Meanwhile, expression levels of inflammatory cytokines in lung tissues were measured by RT-qPCR (Table 6). The left lung lobe from infected mice was fixed in formalin buffer for sectioning, hematoxylin and eosin (H&E) staining, and histopathological examination (Bio-OS Biotechnology Co., Ltd. Wuhan, China).

**Table 6.**
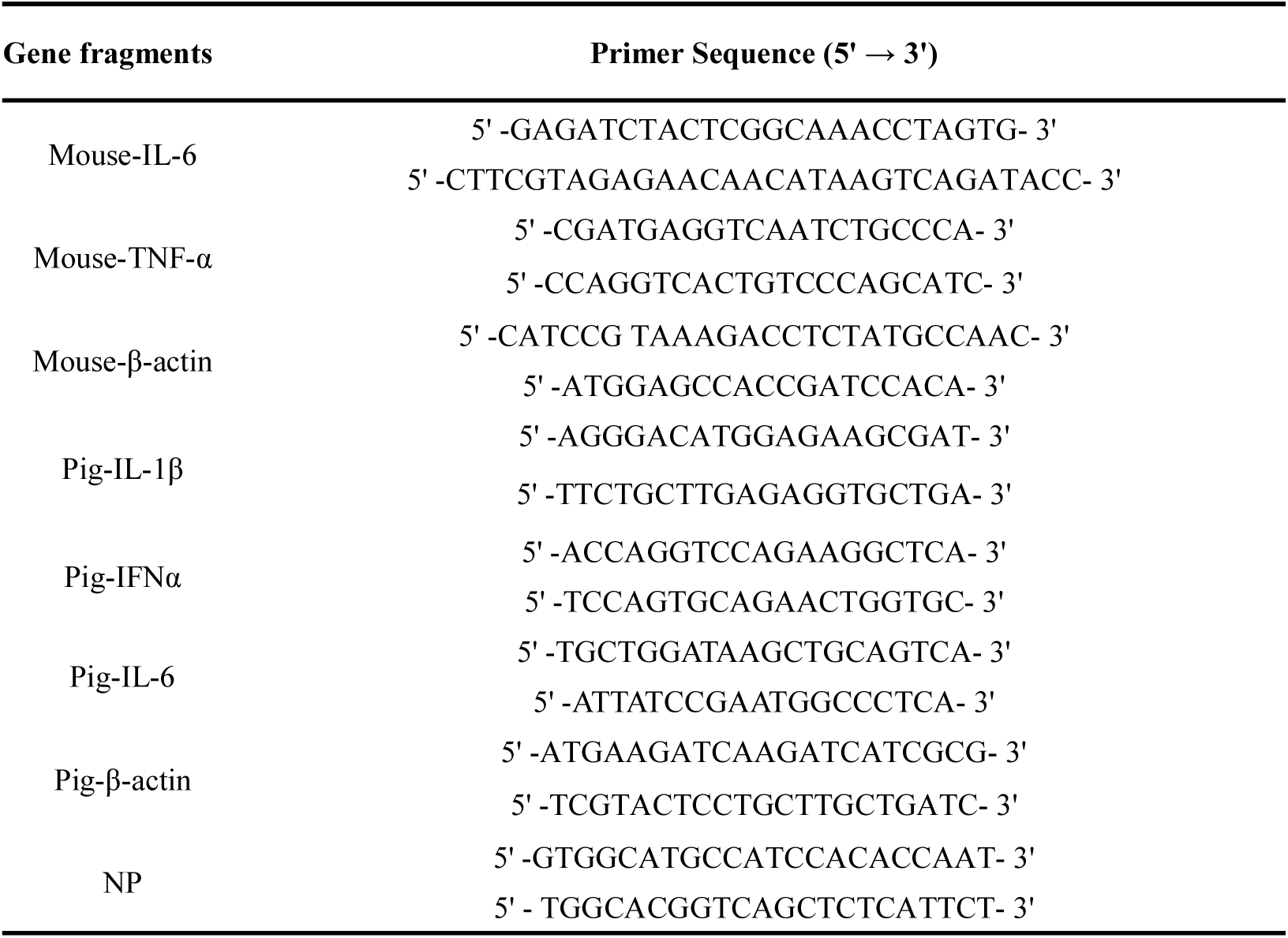
Primers used for dye-based qPCR in this study.

Mice in the weight monitoring and control groups were observed for 12 consecutive days to record mortality, body weight changes, and clinical signs. Weight loss exceeding 25% of the initial body weight was designated as the humane endpoint and recorded as mortality.

### Pathogenicity assessment of two H1N1 SIV strains in pigs

Fifteen 4-week-old piglets seronegative for influenza virus were randomly allocated into three groups (n = 5 per group). Five animals each group were intranasally inoculated with 10^6^ EID_50_ of either the rFYD30 or rYZF69 strain in 1 mL volume, while the control group received 1 mL of PBS via the same route. Body temperature and clinical signs were monitored daily for 14 d. Nasal swabs were collected at 1, 3, 5, 7, 9, and 11 d post-inoculation (dpi) to quantify viral shedding by qPCR. Serum samples were obtained at 7 and 14 dpi for hemagglutination inhibition assay using 1% chicken red blood cells.

On days 3, 5, and 7 post-inoculation, one piglet from each inoculated groups and one from the PBS control group were humanely euthanized. Tissue samples including nasal turbinates, trachea, lung, mandibular lymph nodes, and hilar lymph nodes were collected for viral load quantification, and lung tissue and hilar lymph nodes were measured for inflammatory cytokine levels, both by qPCR (Table 6). Lung lobes from infected and control pigs were fixed in formalin buffer for sectioning, hematoxylin and eosin (H&E) staining, histopathological evaluation, and immunohistochemical staining (Bio-OS Biotechnology Co., Ltd. Wuhan, China). The remaining animals were euthanized on 14 d for final serum collection.

### Cross-hemagglutination inhibition assay

FYD30 and YZF69, together with QD strain and H3N2 SIV previously isolated in the laboratory, were inactivated and used to immunize SPF chickens, and sera were collected for hemagglutination inhibition (HI) assay. The HI assay was performed according to the WOAH (World Organization for Animal Health) standard method, and the HI titer was defined as the highest serum dilution that completely inhibited hemagglutination. The antigenic correlation coefficient (*R*) was calculated as *R =* √𝑟1 × 𝑟2, where *r1* is the ratio of the heterologous serum titer against the heterologous virus to the heterologous serum titer against the homologous virus, and *r2* is the ratio of the homologous serum titer against the heterologous virus to the homologous serum titer against the homologous virus. An *R* value ≥ 0.67 indicated that the two strains belonged to the same antigenic group, whereas an R value < 0.67 indicated a significant antigenic difference.

### Statistical analysis

Statistical analysis was performed in Prism version 8.0.2 (GraphPad Software, San Diego, CA, USA) using the independent-samples *t*-test. For all analyses, *p* < 0.05 (*), *p* < 0.01 (**) was considered as statistically significant between two groups.

## Author contribution statement

All authors contributed to the article and approved the submitted version.

## Data availability statement

The supplementary raw data of this study are all available online: https://doi.org/10.6084/m9.figshare.32181396(50)

## Acknowledgments

The work was partly supported by the National Natural Science Foundation of China (32473040), and A Project Funded by the Priority Academic Program Development of Jiangsu Higher Education Institutions (PAPD). JJ is supported by Postgraduate Research & Practice Innovation Program of Jiangsu Province (SJCX25_2384).

